# A comprehensive resource on chemicals used in aquaculture and their ecotoxicity

**DOI:** 10.64898/2026.01.26.701529

**Authors:** Shreyes Rajan Madgaonkar, Shrish Vashishth, Nikhil Chivukula, Vasavi Garisetti, Shambanagouda Rudragouda Marigoudar, Krishna Venkatarama Sharma, Areejit Samal

**Affiliations:** The Institute of Mathematical Sciences (IMSc), Chennai, India; Homi Bhabha National Institute (HBNI), Mumbai, India; National Centre for Coastal Research, Ministry of Earth Sciences, Government of India, Pallikaranai, Chennai, India

**Keywords:** Aquaculture, Ecotoxicology, Database, Chemical Regulation, One Health, Food Safety

## Abstract

Sustainable aquaculture requires comprehensive chemical oversight, as compounds used in aquaculture can persist in ecosystems, bioaccumulate through food chains, and affect aquatic life and human health. This study presents ReCAnt (<u>Re</u>source on <u>C</u>hemicals used in <u>A</u>quaculture a<u>n</u>d their Ecotoxicity), which compiles information on 690 aquaculture chemicals, with data on toxic effects and therapeutic potential curated from published literature. It was observed that only a fraction of the 690 chemicals are currently regulated, revealing gaps in existing regulations. Integration of data from the Comparative Toxicogenomics Database revealed associations with genes, phenotypes, and diseases, while ECOTOX data provided toxicity and bioconcentration information. Predicted biotransformation pathways and partition coefficients indicated microbial degradation potential and fate across environmental media. Further, food web network analysis identified species vulnerable to trophic transfer and common entry points for chemicals into aquatic ecosystems. This resource can aid in developing evidence-based regulatory frameworks and promoting sustainable chemical management.

## 1. Introduction

Aquaculture has emerged as one of the fastest-growing food production sectors in the world,^1^ surpassing capture fisheries,^2^ and is projected to supply the majority of aquatic dietary protein by 2050,^3^ thereby playing a critical role in addressing ever escalating nutritional demands and food security challenges. Notably, compared to other animal food production systems, aquaculture generates lower average carbon footprints and fewer environmental impacts while providing high nutritional value,^2^ suggesting that the sector will become increasingly important in the shift toward more sustainable food production.

The rapid expansion of aquaculture has intensified reliance on several chemicals to maintain the health of aquatic species and optimize production.^4,5^ Specifically, a broad array of chemicals, including antibiotics, disinfectants, parasiticides, biocides, anesthetics, and immunostimulants,^5^ are intentionally added for their therapeutic effects to combat microbial infections, maintain the health of farmed organisms, and manage water and sediment quality. However, this dependency on intensive chemical use has led to their widespread over-application, posing significant environmental challenges. Despite existing policies, compounds considered legacy contaminants, such as malachite green, which is banned for use in food species due to its documented mammalian toxicity,^6^ continue to persist in aquatic environments,^7^ while approved chemicals such as deltamethrin can lead to mortality in non-target species at very low doses and spread beyond application sites.^8^ These examples highlight significant knowledge and regulatory gaps on the environmental effects of aquaculture chemicals and their potential to cascade through food webs, and eventually affect human health.

Sustainable aquaculture requires addressing the challenges of chemical regulation through a One Health framework.^3,9^ A One Health approach to aquaculture emphasizes the interdependence of farmed organisms, human health, and the environment, and therefore provides a basis for policies and legislation that integrate environmental protection with disease management and production goals.^10–12^ However, current knowledge on the application of aquaculture chemicals, their environmental fate, toxicological profiles, and potential pathways through ecological systems remains fragmented across multiple sources^12^. This gap limits the ability of stakeholders to make informed decisions that balance production efficiency with sustainability and food safety.

To address this need, a comprehensive <u>Re</u>source on <u>C</u>hemicals used in <u>A</u>quaculture a<u>n</u>d their Ecotoxicity (ReCAnt) was developed by consolidating and integrating dispersed data on chemicals used in aquaculture through extensive manual curation of relevant data from published literature and existing toxicological databases. ReCAnt compiles information on physicochemical properties of aquaculture chemicals, their toxicity profiles, predicted environmental fate, and trophic transfer potential through food webs, using both curated data and computational and network-based analyses. In sum, this work presents a novel resource that can support evidence-based evaluation of chemical risks and promote the responsible and sustainable management of chemicals in aquaculture, enabling informed decision-making that helps safeguard the health of aquatic species, humans, and the environment. ReCAnt is accessible at https://cb.imsc.res.in/recant/.

## 2. Methods

### 2.1. Compilation and curation of data on chemicals used in aquaculture

Aquaculture represents one of the fastest-growing food production sectors and relies extensively on chemical interventions, such as antibiotics, algaecides, and parasiticides, to sustain productivity in intensive systems.^13^ In contrast to veterinary drugs like antibiotics,^14^ many other chemicals used in aquaculture, such as biocides,^15^ are subject to limited regulatory oversight, largely because information on their environmental effects remains sparse and fragmented, underscoring the need for a systematic and transparent compilation.

To address this need, a comprehensive list of chemicals used in aquaculture was systematically compiled from published literature through extensive manual curation (**Figure 1**). As a first step, the relevant articles were identified from PubMed (https://pubmed.ncbi.nlm.nih.gov/) using the following query:

> *“aquaculture”[tiab] AND (“drug*”[tiab] OR “chemical*”[tiab])*

This search, last conducted on 21 May 2025, resulted in 2351 articles. Next, these articles were systematically screened in stages to obtain relevant information. First, articles that were not primary research studies or were published in languages other than English were excluded, resulting in 2321 articles. Next, the titles and abstracts of these articles were manually screened for the presence of keywords related to aquaculture or chemical information, resulting in 1457 articles.

**Figure 1:**
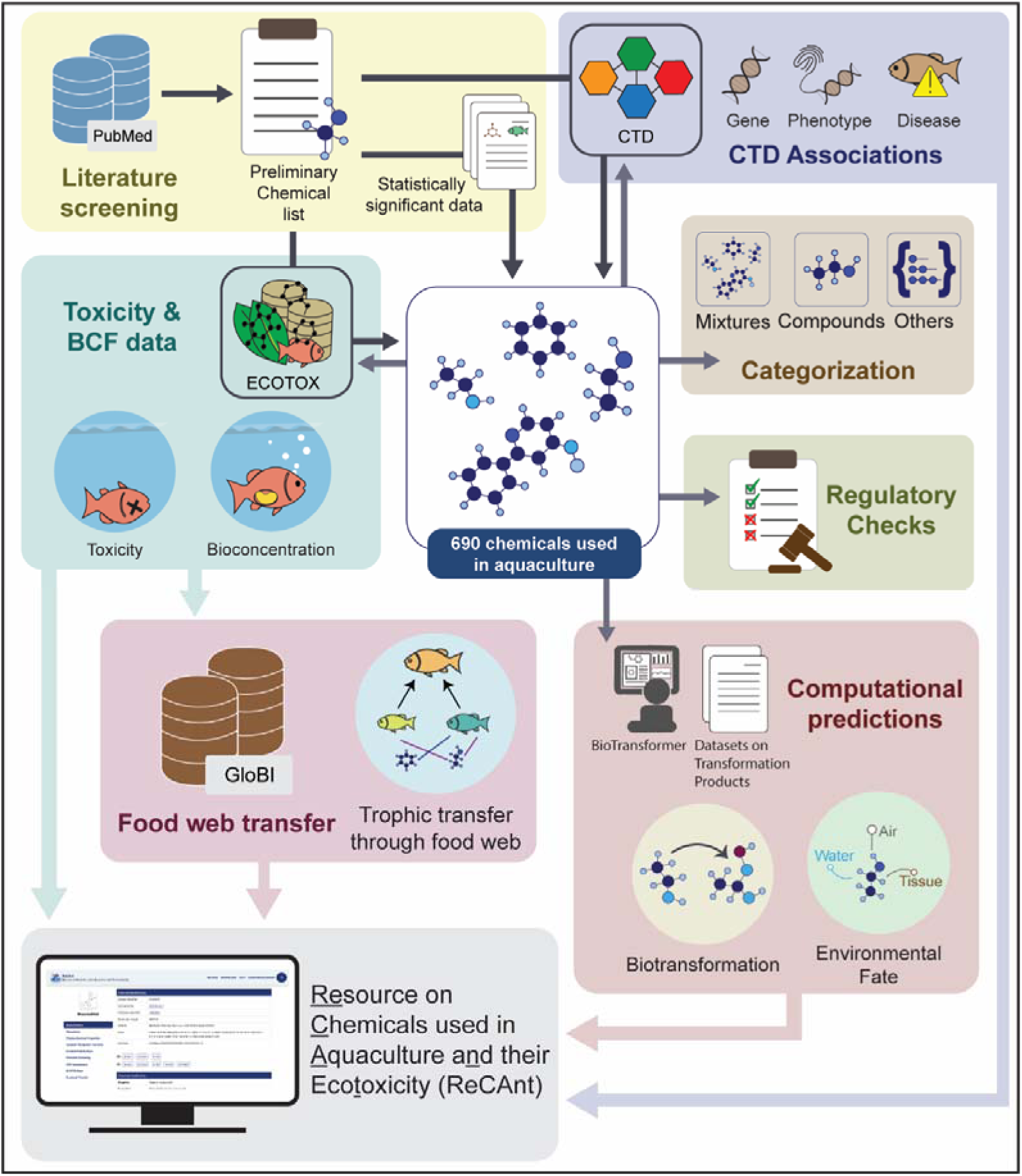
Schematic workflow for the compilation and curation of aquaculture chemicals, their toxicological and therapeutic potential data, and the subsequent development of the ReCAnt (<u>Re</u>source on <u>C</u>hemicals used in <u>A</u>quaculture a<u>n</u>d their Ecotoxicity) webserver.

Thereafter, the full texts of these articles were manually screened for data on therapeutic action or toxic effects of chemicals used in aquaculture, resulting in 257 relevant articles. In addition, information on chemical, chemical type, aquaculture organism in which the chemical was tested or applied, purpose of application, and experimental/field conditions was systematically curated from each of these 257 articles. Subsequently, the chemicals were mapped to corresponding standardized compound identifiers (CIDs) obtained from PubChem (https://pubchem.ncbi.nlm.nih.gov/) and Chemical Abstracts Service registration numbers (CASRN) retrieved from CAS Common Chemistry (https://commonchemistry.cas.org/), resulting in the identification of 748 unique chemicals containing data on therapeutic or toxic effects in at least one research article. The obtained species information was standardized based on genus and species, identifying 249 unique species including 81 aquaculture-relevant species, with the remainder comprising microbes and other pathogens.

Finally, a chemical was considered to be a ‘chemical used in aquaculture’ or ‘aquaculture chemical’ if it contained at least one significant supporting evidence for its therapeutic or toxic effects. Through this extensive manual curation, 682 chemicals were identified with significant evidence from 240 published articles (**Figure 2**). In addition, the associated toxic effects were standardized in terms of Trend, Measurement, and Effect, where these fields represent the observed effect trend relative to controls, the specific experimental endpoint measured, and the overall biological effect, respectively. These definitions were guided by the controlled vocabulary and field definitions used in the ECOTOX^16^ database of the United States Environmental Protection Agency (US EPA), which encodes toxicity results using standardized terms for trend, measurement, and effect to facilitate comparison across studies and organisms.^17^ To ensure comprehensive coverage, the 748 chemicals were further screened in the Comparative Toxicogenomics Database (CTD) and ECOTOX database to identify additional chemicals with relevant data (**Figures 1** and **2**), as described in subsequent sections.

**Figure 2:**
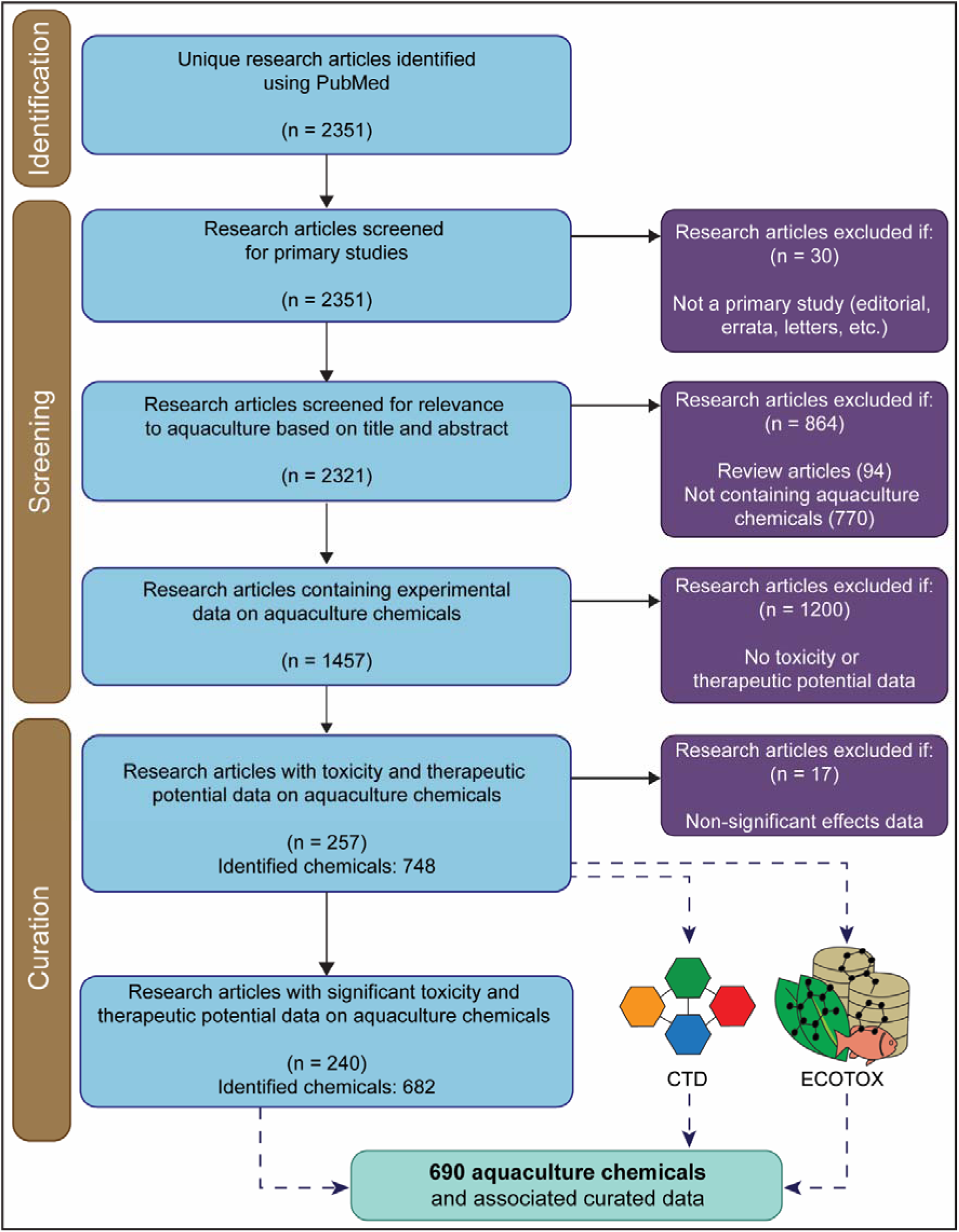
A PRISMA statement outlining the selection of published literature for identifying aquaculture chemicals. Following data extraction and standardization from these studies, additional curation via CTD and ECOTOX resulted in a comprehensive list of 690 chemicals.

### 2.2. Curation of data from Comparative Toxicogenomics Database (CTD)

The Comparative Toxicogenomics Database^18^ (CTD; https://ctdbase.org/) systematically curates associations among chemicals, genes/proteins, phenotypes, and diseases. In addition, CTD provides information on the organism in which each chemical-biological association has been reported, along with the type of association. First, data on chemical-gene, chemical-phenotype, gene-disease, and gene-phenotype associations relevant to aquaculture chemicals were retrieved from the October 2025 release of CTD. Then, the CASRNs of the 748 curated chemicals were used to identify chemical-gene and chemical-phenotype associations within CTD. Further, these associations were filtered to include only data from aquaculture-relevant species.

Next, for chemical-gene associations, gene forms were restricted to mRNA and protein. Further, associations with interaction actions such as ‘secretion’, ‘oxidation’, ‘abundance’, ‘cotreatment’, ‘transport’, ‘reaction’, and ‘response’ were excluded to retain only cases in which the chemical affects gene expression or activity. In the case of chemical-phenotype associations, associations with interaction actions ‘secretion’, ‘oxidation’, ‘abundance’, ‘cotreatment’, ‘transport’, ‘reaction’, ‘response’, ‘metabolic’, and ‘uptake’ were filtered out.

Subsequently, CGPD-tetramers^19^ were constructed using the following criteria established in previous studies^20–22^: (i) chemical-gene and chemical-phenotype associations relevant to aquaculture species and supported by literature evidence; (ii) chemical-disease and gene-disease associations with ‘marker/mechanism’ evidence, and are supported with literature evidence; and (iii) gene-phenotype associations based on Gene Ontology (GO) annotations supported only by experimental evidence (https://geneontology.org/docs/guide-go-evidence-codes/). As gene-disease associations in CTD do not include organism information, only genes obtained from the chemical-gene associations were used to restrict gene-disease associations during CGPD-tetramer construction. Further, these associations were annotated with species information and used to filter out tetramers involving species not relevant to aquaculture. Finally, the resulting chemical-disease associations were manually reviewed to retain only diseases pertinent to aquaculture-relevant species.

### 2.3. Curation of data from ECOTOX

The ECOTOX^16^ database, maintained by the US EPA, is a comprehensive curated repository of ecotoxicological information for over 13,000 chemicals across more than 14,000 ecologically-relevant terrestrial and aquatic species. Here, we relied on ECOTOX to obtain data on chemicals from our list that elicited toxic responses in aquaculture-relevant species as well as showed accumulation within these species. First, the 11 September 2025 version was downloaded via the ‘Download ASCII Data’ option on the ECOTOX website. This dataset comprises information on test organisms, chemical identity, toxicity values, and observed toxicological effects, including biological effects (designated as ‘Effect’), corresponding measurement parameters (designated as ‘Measurement’), and effect trends (designated as ‘Trend’). In addition, the dataset provides bioconcentration factor (BCF) values, which describe the extent of chemical uptake by organisms from their surrounding environment via all exposure routes except dietary absorption.^23^ Relevant information was extracted using an in-house Python script that parsed the tests.txt, results.txt, chemicals.txt, and species.txt files, with chemical records matched using their CASRNs.

In the case of toxicity data, we first restricted to standard acute endpoints, measured as LC50 and EC50, and standard chronic endpoints, measured as NOEL, LOEL, NOEC, and LOEC. Next, the units were normalized to ppm (parts per million) equivalents, and data points having concentration values with units that didn’t allow this conversion were removed. In case of multiple concentration values for a chemical-species pair, the geometric mean of the concentration values was assigned to that pair. Further, data points corresponding to the Trend ‘No Effect’ were removed to retain relevant data. Finally, the species within this data were limited to aquaculture-relevant species.

To understand the uptake of these chemicals, bioconcentration factor (BCF) data were extracted from ECOTOX. First, the units were normalized to L/kg for BCF values, and the species were restricted to aquaculture-relevant species.

Overall, the systematic literature survey and the curation of data from CTD and ECOTOX resulted in the identification of 690 chemicals used in aquaculture, with significant evidence on their therapeutic potential or toxic effects (**Figure 2**; **Tables S1-S3**). These 690 chemicals were then manually classified as ‘compounds’, ‘mixtures’, or ‘others’ based on whether they could be represented as a single, discrete chemical structure suitable for structure-based computation. Plant-derived oils, extracts, multi-component combinations, and commercial formulations were categorized as ‘mixtures’. The remaining chemicals were inspected to assess if they represented single, discrete chemical entities based on available identifiers (CID and CAS), systematic nomenclature, and their association with well-defined chemical classes or functional groups. Chemicals meeting these criteria were categorized as ‘compounds’, while those that did not were assigned to ‘others’. Within the ‘others’ category, chemicals were further subcategorized as ‘nanomaterial’, ‘mineral’, or ‘biological macromolecule’ based on their functional properties and preparation methods described in the literature. Subsequently, these chemicals were assigned unique identifiers and used for further analysis in this study.

### 2.4. Structural, physicochemical, and environmental fate characterization

The two-dimensional (2D) structures for 339 of 690 chemicals were retrieved from PubChem in a structure data file (SDF) format and utilized for downstream structural characterization. Further, Open Babel^24^ was used to generate structural identifiers, derive missing 3D structures by energy minimization, and convert common 2D and 3D structure file formats (**Supplementary Text**). Subsequently, ClassyFire^25^ (http://classyfire.wishartlab.com) was used to classify the chemicals into hierarchical categories, namely, Kingdom, Class, and Subclass, for additional structural annotation. Physicochemical properties were computed using the RDKit module in Python, while 2D and 3D molecular descriptors were calculated using PaDEL^26^ (http://www.yapcwsoft.com/dd/padeldescriptor/), RDKit, and Pybel^27^ (**Supplementary Text**).

The aquaculture chemical space was visualized via a chemical similarity network (CSN)^28^ constructed based on pairwise chemical similarity values computed using the Tanimoto coefficient^29^ based on ECFP4 fingerprint^30^ (**Supplementary Text**). In addition, Bemis–Murcko scaffolds^31^ were generated and visualized using a scaffold cloud representation^32^ to examine scaffold-level similarity across the aquaculture chemical space (**Supplementary Text**).

To understand the regulatory landscape of aquaculture chemicals, lists comprising substances of concern were compiled from aquaculture-relevant regulations provided by regulatory bodies across various jurisdictions (**Supplementary Text**). In addition, persistence, bioaccumulation, mobility, and toxicity (PBMT) assessments were compiled from the European Chemical Agency’s (ECHA) persistence, bioaccumulation, and toxicity (PBT) assessment list (https://echa.europa.eu/pbt) and relevant datasets available via the NORMAN Suspect List Exchange (SLE; https://www.norman-network.com/?q=suspect-list-exchange).

To characterize the fate of the aquaculture chemicals in the environment, their biotransformation products, partitioning behavior, and aqueous solubility were assessed. Transformation products (TPs) were obtained using the BioTransformer 3.0 tool^33^ and curated datasets from the NORMAN SLE. Subsequently, the underlying transformation reactions were represented as a reaction network, which was analyzed using the NetworkX^34^ module and visualized in Cytoscape version 3.10^35^ (**Supplementary Text**). Thereafter, three widely used partition coefficients, namely K_ow_ (octanol-water coefficient), K_aw_ (air-water coefficient), and K_oa_ (octanol-air coefficient), were predicted using the US EPA Estimation Programs Interface (EPI) Suite (https://episuite.dev/EpiWebSuite/<u>#/</u>) (**Supplementary Text**). These partition coefficients were then compared for the aquaculture chemicals to inform predictions of their environmental distribution and dominant partitioning behavior. Finally, the solubility of aquaculture chemicals was interpreted based on aqueous solubility data retrieved from the AqSolDB^36^ (**Supplementary Text**).

### 2.5. Assessment of the trophic transfer potential of aquaculture chemicals

To investigate the potential for trophic transfer of aquaculture chemicals, food-web interaction data from the Global Biotic Interactions (GloBI) database^37^ (https://www.globalbioticinteractions.org/) were integrated with prey-level chemical toxicity and BCF information obtained from ECOTOX. Briefly, interactions in which aquaculture-relevant species function as predators were retrieved from the GloBI database, and thereafter, BCF, acute toxicity, and chronic toxicity data were obtained from ECOTOX for the associated prey species (**Supplementary Text**). This integration enabled the construction of directed networks in which edge weights represent toxicity concentrations or BCF values. Further, the networks were analyzed using NetworkX^34^ to identify aquaculture-relevant species potentially impacted by chemical exposure at lower trophic levels, and the networks were visualized in Cytoscape version 3.10^35^.

## 3. Results and Discussion

### 3.1. Overview of aquaculture chemicals within ReCAnt

The aquaculture sector is one of the largest providers of aquatic protein, becoming an important component of world food security. Globally, over 500 aquatic species are farmed in this sector for different purposes.^3^ However, the intensive nature of modern aquaculture systems necessitates the use of various chemicals, including antibiotics, disinfectants, growth promoters, anesthetics, and therapeutic agents, to maintain the health of aquatic species, prevent disease outbreaks, and optimize production.^11,13^ Therefore, understanding the fate, behavior, and effects of these chemicals is essential for ensuring environmental sustainability and the safety of aquaculture products.

As the sector continues to expand, there is an increasing interest from the scientific research community in such aquaculture chemicals. The publication trend for such research shows steady growth, with papers nearly doubling every five years (**Figure 3A**). Therefore, this study presents a comprehensive resource, the <u>Re</u>source on <u>C</u>hemicals used in <u>A</u>quaculture a<u>n</u>d their Ecotoxicity (ReCAnt), that has compiled and curated data from existing literature on aquaculture chemicals to bridge the gap between fragmented information sources and the growing need for systematic, accessible data on chemical properties, regulatory coverage, environmental fate, and biological effects in aquaculture systems.

**Figure 3:**
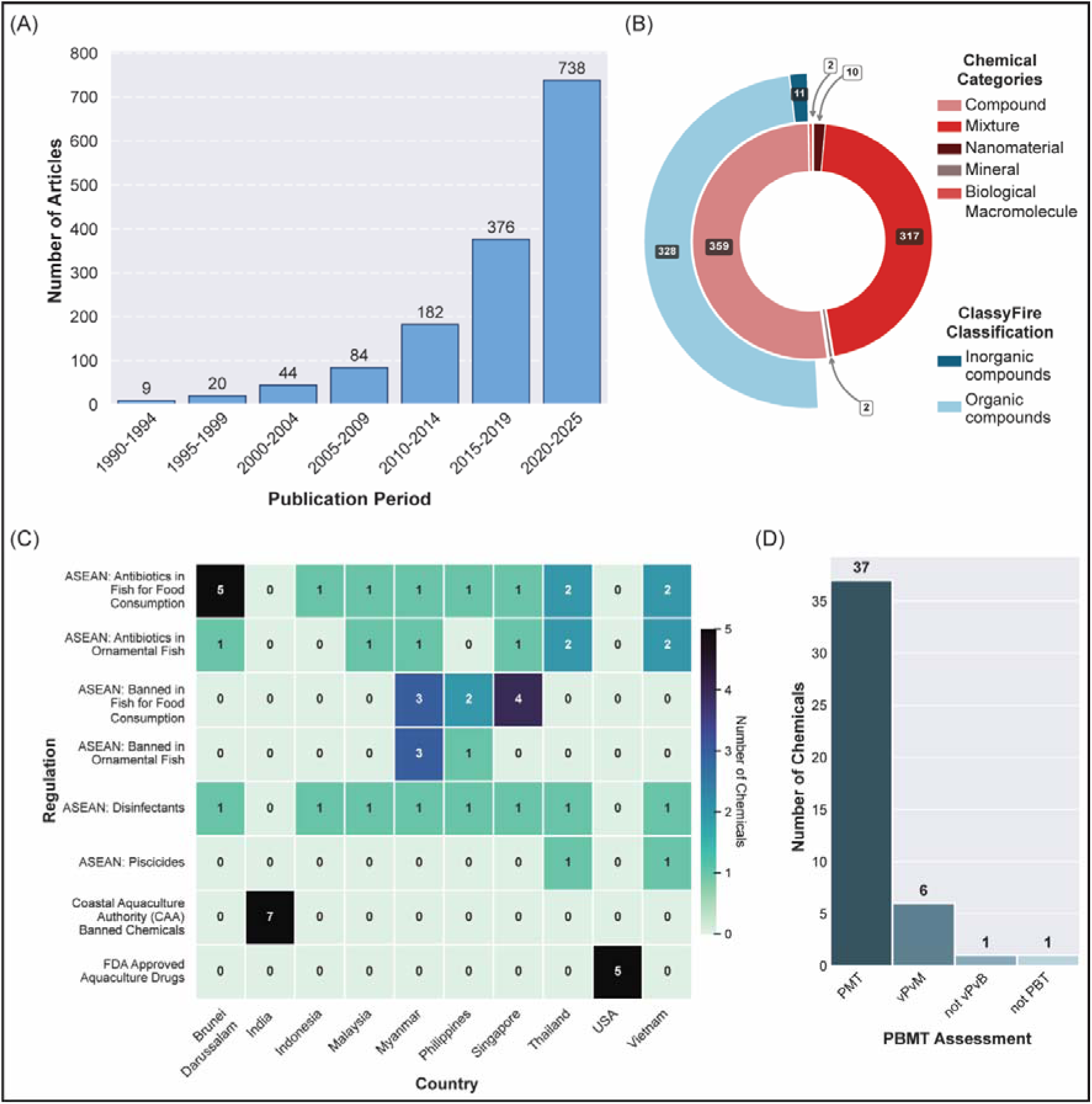
**(A)** Publication trend for aquaculture chemical research from 1990 to 2025. **(B)** Sunburst plot depicting the distribution of aquaculture chemicals across different categories within ReCAnt (<u>Re</u>source on <u>C</u>hemicals used in <u>A</u>quaculture a<u>n</u>d their Ecotoxicity). The inner ring distinguishes chemicals by the discreteness of their molecular structure, while the outer ring provides a ClassyFire-based classification of 339 discrete compounds into organic and inorganic categories. **(C)** Heatmap depicting the presence of aquaculture chemicals in chemical regulations across different countries. The number of aquaculture chemicals within each of the chemical regulations is denoted in the heatmap. **(D)** Distribution of aquaculture chemicals across Persistence, Bioaccumulation, Mobility, and Toxicity (PBMT) assessment categories.

ReCAnt primarily provides information on 690 chemicals (**Table S1**) used in aquaculture, covering their toxic effects (**Table S2**) and therapeutic potential (**Table S3**). Toxic effects data were available for 215 chemicals and therapeutic potential data for 621 chemicals, with mortality being the most represented toxic effect and antibacterial activity the most commonly reported therapeutic action. The resource comprises 359 compounds, 317 mixtures, and 14 other chemical entities, including nanomaterials, biological macromolecules, and minerals (**Figure 3B**; **Table S1**). Notably, the majority of mixtures are plant-derived natural products or essential oils, suggesting a shift toward natural alternatives in aquaculture practices. Moreover, ReCAnt compiles chemical structures of 339 compounds, of which 328 are organic while the remaining 11 are inorganic (**Figure 3B**; **Table S1**). Combined with the predominance of plant-derived organics among the mixtures, this indicates that the chemicals in our database are primarily organic in nature.

### 3.2. Integrated Chemical Data from CTD and ECOTOX

The Comparative Toxicogenomics Database^18^ (CTD) provided associations for 54 chemicals with 7997 genes (**Table S4**) and 49 chemicals with 231 phenotypes (**Table S5**). CGPD-tetramer construction integrated these data to generate 768 tetramers comprising 17 chemicals, 79 genes, 48 phenotypes, and 125 diseases. Following filtering for aquaculture-relevant species and manual curation, chemical-disease associations were identified for 17 chemicals with 89 diseases (**Table S6**). Atrazine (CID:2256) exhibited the highest number of gene associations at 4866, while chlorpyrifos (CID:2730) showed the most associations with phenotypes at 96 and diseases at 57. Among disease associations, ‘Chemical and Drug Induced Liver Injury’ was most frequently represented, followed by ‘Acute Kidney Injury’ and ‘Kidney Diseases’.

ECOTOX^16^ contained toxicity data for 142 chemicals tested across 53 aquaculture-relevant species. Acute toxicity endpoints (LC50, EC50) were available for 119 chemicals across 48 species (**Table S7**), while chronic toxicity endpoints (NOEC, NOEL, LOEL) are available for 112 chemicals across 49 species (**Table S7**). The toxicity concentration distributions were right-skewed, with median values of 2.30 ppm equivalents for acute toxicity and 0.48 ppm equivalents for chronic toxicity. Copper sulfate (CID:24462) had the most extensive data records for both toxicity types, followed by sodium hypochlorite (CID:23665760) for acute toxicity and atrazine (CID:2256) for chronic toxicity. Additionally, bioconcentration factor (BCF) data were available for 24 chemicals across 16 species (**Table S8**). The BCF distribution showed a median of 86.74 L/kg.

### 3.3. Regulatory coverage of aquaculture chemicals within ReCAnt

Several aquaculture-relevant regulations were relied upon to gauge the regulatory coverage of the chemicals compiled in ReCAnt. It was observed that 7 chemicals within ReCAnt were among the 29 antibiotics banned for shrimp aquaculture in India^38^ (**Figure 3C**; **Table S9**). Moreover, 5 chemicals were identified by the US FDA as approved drugs for aquaculture (**Figure 3C**; **Table S9**). It was observed that 27 chemicals were cataloged in 10 of the 12 chemical monitoring lists across 8 ASEAN countries^39^ (**Figure 3C**; **Table S9**). Among these 27 chemicals, 6 chemicals were banned from usage in Brunei Darussalam, 4 were banned in Singapore, Thailand and Vietnam, 3 were banned in Myanmar, and 2 were banned in Indonesia, Malaysia, and Philippines. These banned chemicals are categorized across 6 regulatory lists, namely Antibiotics in Fish for Food Consumption (5 chemicals), Banned in Fish for Food Consumption (4 chemicals), Banned in Ornamental Fish (3 chemicals), Antibiotics in Ornamental Fish (2 chemicals), Disinfectants (1 chemical), and Piscicides (1 chemical) (**Table S9**). Furthermore, 39 chemicals were identified as substances of concern across different regulatory lists (**Figure S1**; **Table S9**). Moreover, 45 and 48 chemicals were found in the OECD HPV list and the US HPV list, respectively, with a combined total of 50 chemicals produced in excess of one ton per year across the US and OECD regions (**Figure S1**; **Table S9**).

Further, only one compound, nerolidol (CID:5284507), was found in the ECHA REACH PBT list with assessments of ‘not PBT’ and ‘not vPvB’ (**Table S9**). The datasets UBAPMT,^40^ EAWAGPMT,^41^ and UFZHSFPMT^42^ provided assessments for 6, 25, and 12 chemicals, respectively (**Table S9**). Overall, across the three datasets, 37 compounds had PBMT assessments, with all compounds assessed as PMT (**Figure 3D**). Notably, six chemicals, namely azithromycin (CID:447043), sulfadiazine (CID:5215), erythromycin (CID:12560), atrazine (CID:2256), triclosan (CID:5564), and diuron (CID:3120) were additionally assessed as vPvM (**Figure 3D**). This indicates that these compounds are both very persistent and very mobile in the environment.

### 3.4. Chemical space of aquaculture chemicals

The chemical similarity network (CSN) consisted of 339 nodes and 148 edges, retaining only those edges with a Tanimoto coefficient greater than 0.5 (**Figure S2**; **Table S10**). Among the 229 components in the network, 53 include two or more nodes. The largest connected component (LCC) consists of 13 nodes and 20 edges. Further, 32 of the 53 components consisted of 2 nodes, suggesting that the network is highly fragmented into smaller isolated clusters. The degree distribution showed that 176 nodes were isolated nodes, while 96 nodes had a degree of 1, followed by 35 nodes with a degree of 2. The network’s disconnectedness was further highlighted by the average degree and average betweenness centrality, which were 1.816 and 0.0001, respectively.

The scaffold cloud representation (**Figure 4**) depicts 35 chemical scaffolds derived from at least two compounds (out of 184 unique scaffolds identified), with quinone being the most represented scaffold, followed by benzene, with frequencies of 26 and 24, respectively. The next highest scaffold frequency was 6, suggesting that the 339 aquaculture chemicals are structurally diverse beyond these two dominant scaffolds.

**Figure 4:**
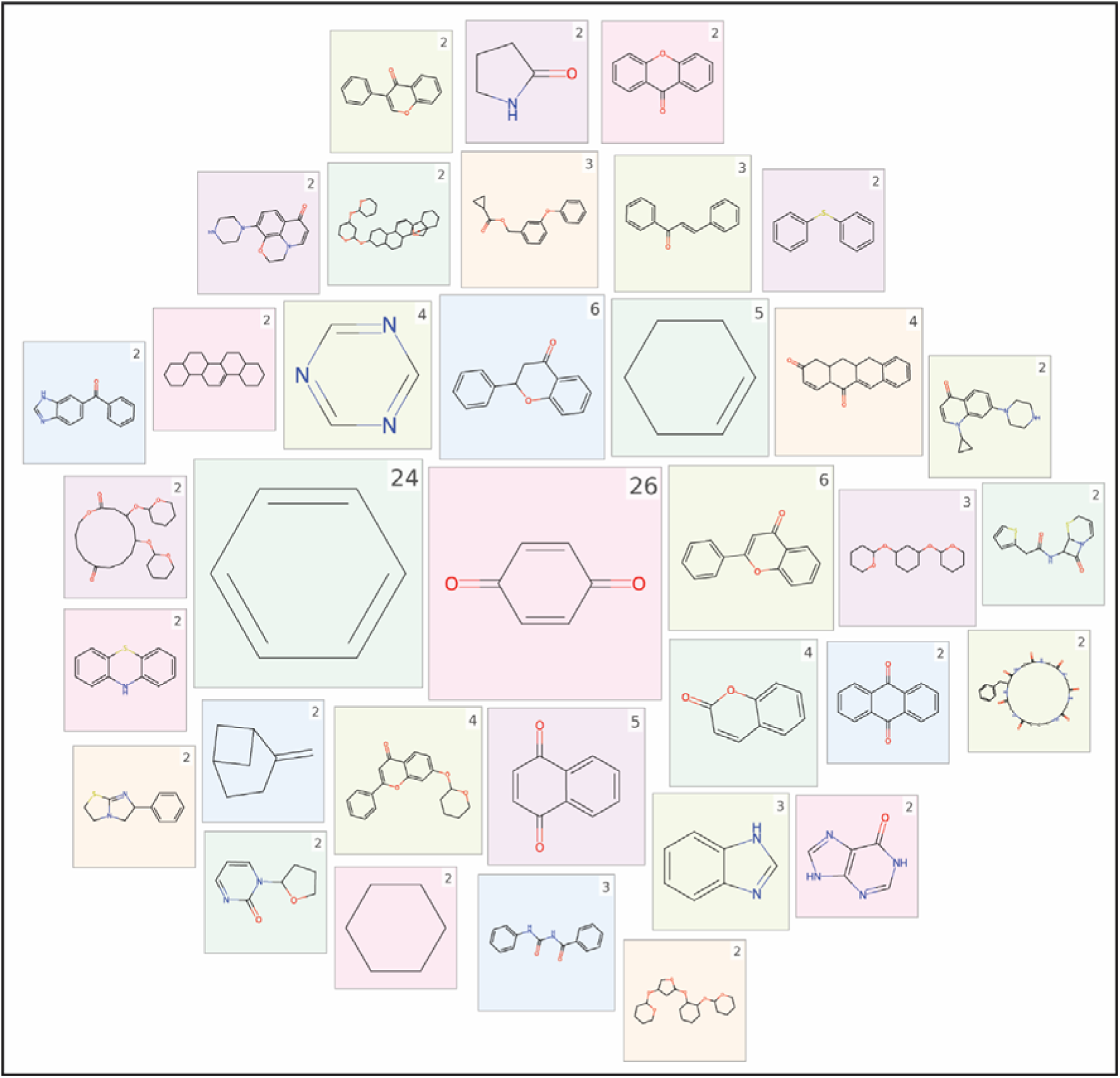
A scaffold cloud representation of the 35 unique scaffolds shared by at least two chemicals, identified from 339 discrete compounds in ReCAnt. Scaffolds were computed using RDKit based on the Bemis-Murcko definition. The top-right corner of each tile denotes the number of chemicals sharing that specific scaffold.

The high fragmentation of the CSN, combined with the scaffold frequency distribution, indicates that aquaculture chemicals exhibit considerable structural diversity with limited chemical similarity among most compounds. It should be noted that mixtures and other chemical entities lacking discrete structural representations were excluded from this analysis.

### 3.5. Biotransformation and environmental fate

BioTransformer 3.0^33^ successfully predicted transformation products (TPs) for 228 chemicals, while the REFTPS^43^ dataset provided TPs for 11 chemicals. Overall, biotransformation data were compiled for 229 unique chemicals of the 339 aquaculture chemicals with structural data (**Table S11**). The directed network for these biotransformation reactions comprised 1380 edges connecting 1510 nodes, of which 1281 nodes corresponded to TPs (**Figure S3**). As no overlap was observed between aquaculture chemicals and their predicted TPs, the network exhibited a hierarchical, acyclic structure. Atrazine (CID:2256) showed an out-degree of 39, which is the highest among all aquaculture chemicals, followed by ivermectin (CID:6321424) and netilmicin (CID:441306) with out-degrees of 25 and 22, respectively. Among TPs, acetaldehyde (CID:177) and methanol (CID:887) showed the highest in-degree of 10, followed by formate (CID:283) with an in-degree of 7. This potentially suggests that several complex molecules eventually converge into the same few simple molecules.

The directed network consisted of 159 weakly connected components, with the largest weakly connected component comprising 403 nodes and 418 edges. This component included 46 aquaculture chemicals and 357 TPs and comprised nodes with the highest in- and out-degrees identified above.

Partition coefficients were predicted for 230 aquaculture chemicals with available structural information using the US EPA EPI Suite (**Table S1**). Following the approach defined by Mackay and Parnis,^44,45^ the propensity of a chemical to partition into air, water, or octanol was examined by plotting logK_aw_ against logK_ow_, with the logK_oa_ values representing a color scale (**Figure 5A**). It was observed that most chemicals were located in the upper-left region of the plot, indicating relatively higher air-water partitioning compared to octanol-water partitioning, consistent with higher volatility.^44^ Similarly, when predicted BCF and bioaccumulation factor (BAF) were examined as color scales (**Figures 5B, C**), a low proportion of chemicals showed higher bioconcentration and bioaccumulation potential.

**Figure 5:**
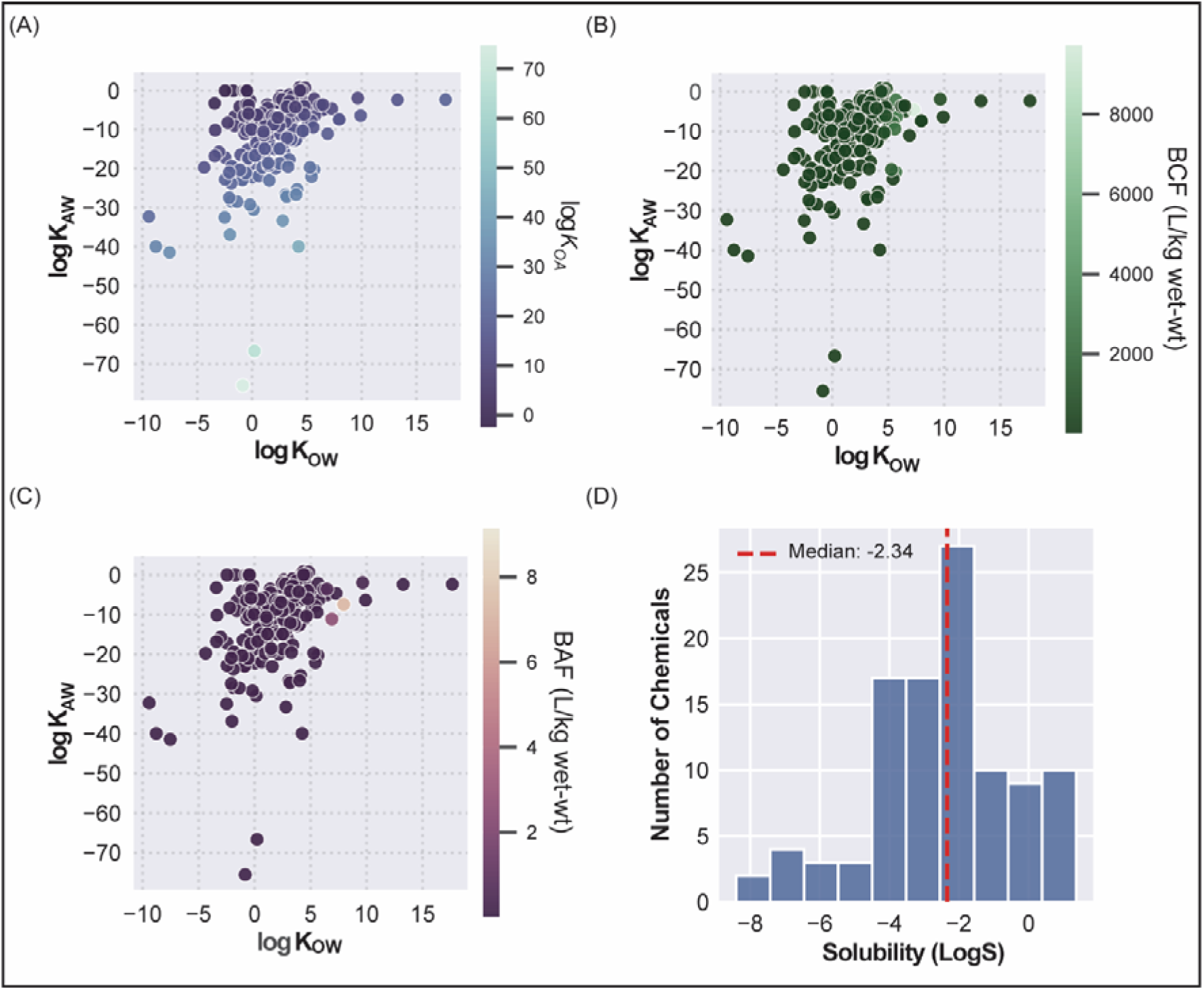
Plots of air-water coefficient (logK_aw_) against octanol-water coefficient (logK_ow_), with the **(A)** octanol-air coefficient (logK_oa_) values, **(B)** Bioconcentration Factor (BCF), and **(C)** Bioaccumulation Factor (BAF) represented on a color scale. **(D)** Distribution of aqueous solubility (LogS) values for 102 of the 339 discrete aquaculture chemicals.

Further, the aqueous solubility was assessed for 102 of the 339 aquaculture chemicals that had solubility data from AqSolDB^36^ (**Table S1**). It was noted that the LogS values spanned a wide range from −8.402 to 1.365, indicating large variability in aqueous solubility. The median LogS (−2.34) falls within the slightly soluble category, suggesting that at least half of the compounds in the dataset exhibit low aqueous solubility, with a considerable proportion approaching insolubility (**Figure 5D**).

Overall, these findings indicate that aquaculture chemicals generally exhibit low to moderate aqueous solubility. Moreover, the partitioning hierarchy indicates that many chemicals favor atmospheric partitioning over partitioning into aqueous or organic phases.

### 3.6. Food web-based trophic transfer

In this study, dietary exposure of aquaculture-relevant species to the compiled aquaculture chemicals was assessed by constructing food web networks. In these networks, the prey organisms of the target species were integrated with toxicity and BCF data for aquaculture chemicals to identify pathways through which chemicals could cause secondary toxic effects or bioaccumulate in aquaculture-relevant species.

First, a food web network was constructed using BCF values (**Figure S4**) to establish links between chemicals and prey species. This resulted in a network comprising 348 edges (**Table S12**) and 120 nodes (**Table S13**), of which 24 were chemicals, 47 were prey species, 39 were predators, and 10 species functioned as both predator and prey. For each chemical-predator pair, path existence served as a proxy for bioaccumulation potential through trophic transfer. Irgarol (CID:91590) is connected to 40 predator species, followed by atrazine (CID:2256) and chlorpyrifos (CID:2730), connected to 31 and 27 species, respectively. Among predators, *Micropterus salmoides* was connected to 28 chemicals, suggesting potential vulnerability to bioaccumulation through dietary intake, followed by *Salmo trutta* to 27 chemicals and *Oncorhynchus mykiss* to 23 chemicals.

Next, food web networks were constructed using acute and chronic toxicity data. The network based on acute toxicity data comprised 2460 edges (**Table S12**) and 440 nodes (**Table S13**), of which 127 are chemicals, 244 are prey species, 38 are predators, and 31 are species that are both predators and prey. The network based on chronic toxicity data comprised 2365 edges (**Table S12**) and 434 nodes (**Table S13**), of which 113 are chemicals, 281 are prey species, 40 are predators, and 30 are species that are both predators and prey. In the acute toxicity-based food web network, dichlorvos (CID:3039) was connected to 62 predators, followed by copper sulfate (CID:24462) and cypermethrin (CID:2912), connected to 59 and 58 predator species, respectively. In the chronic toxicity-based food web network, chlorpyrifos (CID:2730) was connected to 67 predators, followed by malathion (CID:4004) and simazine (CID:2516), connected to 64 and 62 predator species, respectively. *Micropterus salmoides* had the most chemical connections, with 127 chemicals in the acute toxicity-based network and 100 in the chronic toxicity-based network.

Finally, betweenness centrality was calculated for prey species to identify those appearing most frequently on paths connecting chemicals to predators. This calculation treated the network as unweighted, as connections between predator and prey species carried an edge weight of 0. Species with the highest betweenness centrality values signify the most frequent points of dietary exposure to aquaculture chemicals. *Gammarus pulex* had the highest betweenness centrality in the food web constructed based on BCF values, while *Raphidocelis subcapitata* and the class Insecta were most central in the acute and chronic toxicity networks, respectively (**Table S13**).

### 3.7. Features of the ReCAnt webserver

The compiled data and visualizations generated in this study for 690 aquaculture chemicals are available through a dedicated online resource, namely, the <u>Re</u>source on <u>C</u>hemicals used in <u>A</u>quaculture a<u>n</u>d their Ecotoxicity (ReCAnt), accessible at: https://cb.imsc.res.in/recant/. For each chemical, ReCAnt provides comprehensive information, including molecular descriptors, physicochemical properties, therapeutic potential, toxic effects, CTD associations (genes, phenotypes, diseases), and ECOTOX-derived toxicity and bioconcentration data. The resource integrates predicted biotransformation products with network-based visualizations depicting transformation pathways, and provides predator-centric food web visualizations highlighting potential trophic transfer routes. All database identifiers are hyperlinked to their respective sources: chemical identifiers (CID, CAS) link to PubChem (https://pubchem.ncbi.nlm.nih.gov) and Common Chemistry CAS (https://commonchemistry.cas.org/), CTD genes to NCBI Gene (https://www.ncbi.nlm.nih.gov/gene/), phenotypes (GO terms) to AmiGO Gene Ontology (https://amigo.geneontology.org/), diseases (MeSH terms) to NCBI MeSH (https://meshb.nlm.nih.gov/), and PubMed IDs to PubMed articles (https://pubmed.ncbi.nlm.nih.gov/). Additional details on the technical implementation of the ReCAnt web interface are provided in the **Supplementary Text**.

## 4. Conclusion

Sustainable aquaculture requires integrated chemical oversight as chemicals used can persist in ecosystems, bioaccumulate through food chains, and subsequently endanger aquatic life, ecosystems, and human health. In this study, a comprehensive resource, ReCAnt (<u>Re</u>source on <u>C</u>hemicals used in <u>A</u>quaculture a<u>n</u>d their Ecotoxicity), is presented, compiling information on 690 aquaculture chemicals, with data on toxic effects and therapeutic potential curated from published literature for 215 and 621 chemicals, respectively. Notably, only a fraction of the curated chemicals are currently regulated, with several classified as persistent, mobile, and toxic (PMT) substances, highlighting critical gaps in existing regulatory frameworks. Integration of data from the Comparative Toxicogenomics Database (CTD) for 63 aquaculture chemicals revealed associations spanning 7997 genes, 231 phenotypes, and 89 diseases, with varying data coverage across chemicals. In addition, biotransformation products and partition coefficients were predicted for 228 and 230 chemicals, respectively. Analyses indicated that these chemicals potentially undergo biotransformation into simpler compounds through microbial degradation, and can readily partition into atmospheric media. Moreover, toxicity and bioconcentration factor (BCF) data was obtained from ECOTOX for 142 and 24 chemicals, respectively. Integration of food web networks with ECOTOX data enabled the identification of aquaculture-relevant species most vulnerable to trophic transfer through dietary exposure, as well as prey species representing frequent entry points for aquaculture chemicals into food webs. Together, these integrated analyses characterize aquaculture chemicals across toxicological effects, environmental behavior, and trophic transfer through food webs. All data generated in this study are publicly available through the online resource for academic research and linked to other resources for enhanced interoperability.

While ReCAnt provides a comprehensive assessment of aquaculture chemical impacts, several factors limit the scope of the study. Data curation was limited to peer-reviewed research articles indexed in PubMed, excluding regulatory agency reports, industry documents, and other non-indexed sources that may contain additional relevant information. Additionally, standardized structural information was available only for approximately half of the curated chemicals, as mixtures and other chemical entities lacked chemical identifiers and frameworks for systematic integration with external, restricting these chemicals to manually curated toxicity and therapeutic data only. Further, biotransformation and environmental fate predictions rely on computational tools that may not fully capture the behavior of these chemicals under natural environmental conditions. Moreover, the list of aquaculture-relevant species used in this study, while extensive, is not exhaustive and may not cover all farmed species across diverse aquaculture systems.

Nonetheless, ReCAnt is the first resource that extensively compiles chemicals used in aquaculture. It provides data on their structural and physicochemical characteristics and integrates information from toxicological databases, making this information more accessible to researchers, regulators, and other stakeholders. Furthermore, the resource utilizes computational tools and network-based methods to assess biotransformation pathways, partitioning behavior, and trophic transfer potential within aquaculture-relevant environments. In the future, the resource can be further developed to expand the chemical coverage, integrate additional biologically relevant data, explore frameworks for mixture toxicity assessment, and assess pathogen adaptation to aquaculture chemicals. In conclusion, this study presents a novel resource on aquaculture chemicals that can aid in developing evidence-based regulatory frameworks and promoting sustainable chemical management aligned with One Health principles, safeguarding aquatic ecosystems, food security, and human health.

## Acknowledgement

The authors would like to thank Dr. K. Ramu for discussions. Areejit Samal would like to acknowledge funding from the Department of Atomic Energy (DAE), Government of India via Apex project to The Institute of Mathematical Sciences (IMSc) Chennai. This work was also undertaken as part of the project on ‘Marine Ecotoxicology and Ecological Risk Assessment’ (MEERA) Programme [MoES/OSMART/EFC/2021 dated 07.03.2022] at the National Centre for Coastal Research (NCCR), Ministry of Earth Sciences, Government of India.

## CRediT author contribution statement

**Shreyes Rajan Madgaonkar:** Conceptualization, Data Curation, Formal Analysis, Methodology, Visualization, Writing; **Shrish Vashishth:** Conceptualization, Data Curation, Methodology, Writing; **Nikhil Chivukula:** Formal Analysis, Visualization, Writing**; Vasavi Garisetti:** Data Curation; **Shambanagouda Rudragouda Marigoudar:** Conceptualization, Formal Analysis, Methodology, Writing; **Krishna Venkatarama Sharma:** Conceptualization, Formal Analysis, Methodology, Writing; **Areejit Samal:** Conceptualization, Supervision, Formal Analysis, Methodology, Writing.

## Supplementary Tables

**Table S1:** This table contains information on the curated list of 690 chemicals used in aquaculture. For each chemical, the table provides the chemical name, PubChem Identifier, Chemical Abstracts Service Registry Number (CASRN), category, molecular weight (in g/mol), canonical SMILES, InChI, InChIKey, ClassyFire classifications (Kingdom, SuperClass, and Class), the DSSTox identifier, the DSSTox IUPAC name, and the DSSTox molecular formula curated from the US EPA’s CompTox Chemicals Dashboard. Additionally, for each chemical, the table provides physicochemical and bioaccumulation properties, including predicted and experimental log□Kow values, predicted log□Kaw value, predicted and experimental log□Koa values, bioconcentration factor (BCF) value, bioaccumulation factor (BAF) value, and aqueous solubility value and associated standard deviation (in terms of LogS). Predicted partition coefficients (log□Kow, log□Kaw, log□Koa), BCF, and BAF are obtained from EPA EPI Suite, and solubility data is obtained from AqSolDB.

**Table S2:** This table contains information on the toxic effects of 215 aquaculture chemicals. For each chemical, the table provides the chemical name, PubChem Identifier, Chemical Abstracts Service Registry Number (CASRN), the chemical category, the PubMed literature identifier of corresponding evidence (PMID), the type of experiment, the cell or tissue used, the aquaculture-relevant test organism, the life stage of that organism, the method of administration, the unit of the administered or significant dose(s), the value of administered and significant dose(s), and the standardized toxicity endpoints (trend, measurement, and effect).

**Table S3:** This table contains information on the therapeutic potential of 621 aquaculture chemicals. For each chemical, the table provides the chemical name, PubChem identifier, Chemical Abstracts Service Registry Number (CASRN), the chemical category, the PubMed literature identifier of corresponding evidence (PMID), the type of experiment, the cell, tissue, or microbial strain used, the aquaculture-relevant test organism, the life stage of that organism, the target organism within the host, the method of administration, the unit of the administered or significant dose(s), the administered and significant dose(s), the time to death of the target organism (in minutes), the anthelminthic efficacy of the chemical (in %), and the therapeutic action.

**Table S4:** This table contains information on the CTD chemical-gene associations for 54 chemicals and 7997 genes. For each association, the table provides the PubChem identifier and Chemical Abstracts Service Registry Number (CASRN) of the chemical, the gene form of the associated gene, the NCBI gene identifer, the gene symbol, the interaction action between chemical and gene, the NCBI taxonomy identifier and the Latin name of associated organism, and the literature identifier of corresponding reference(s) (separated by ‘|’ symbol).

**Table S5:** This table contains information on CTD chemical-phenotype associations for 49 chemicals and 239 phenotypes. For each association, the table provides the PubChem identifier and Chemical Abstracts Service Registry Number (CASRN) of the chemical, the associated phenotype identifier from gene ontology (GO), the phenotype name, the interaction action between chemical and phenotype, the NCBI taxonomy identifier and the Latin name of associated organism, and the literature identifier of corresponding reference(s) (separated by ‘|’ symbol).

**Table S6:** This table contains information on the CTD chemical-disease associations for 17 chemicals and 89 diseases identified through Chemical-Gene-Phenotype-Disease (CGPD) tetramer construction. For each association, the table provides the PubChem identifier and Chemical Abstracts Service Registry Number (CASRN) of the chemical, the Medical Subjects Headings (MeSH) identifier of the disease, the disease name, and the Latin name of the associated organism(s) (separated by ‘|’ symbol).

**Table S7:** This table contains information on 142 aquaculture chemicals associated acute and chronic toxicity endpoint data from ECOTOX. For each chemical, the table provides the PubChem identifier, Chemical Abstracts Service Registry Number (CASRN), the Latin name of the associated species, the corresponding ECOTOX species group, the NCBI taxonomy identifer for the species, the type of toxicity, the toxicity endpoint, the toxicity concentration (in ppm equivalents), and the trend, measurement, and effect associated with the observed effects.

**Table S8:** This table contains information on bioconcentration factor (BCF) data from ECOTOX associated with 24 aquaculture chemicals. For each chemical, the table provides the PubChem identifier, Chemical Abstracts Service Registry Number (CASRN), the Latin name of the associated species, the corresponding ECOTOX species group, the NCBI taxonomy identifier for the species, and the BCF value (in L/kg).

**Table S9:** This table contains data on the aquaculture chemicals present across aquaculture-relevant regulations, substances of concern lists, substances in use lists, and high production volume (HPV) chemicals lists, along with their Persistence, Bioaccumulation, Mobility, and Toxicity (PBMT) assessment. For each chemical, the table provides the chemical name, PubChem identifier, Chemical Abstracts Service Registry Number (CASRN), the name of the aquaculture-specific regulatory list, the associated country and regulatory status, the name, class, and link to the corresponding source data file for other regulatory lists. For the PBMT assessment data, the table provides the corresponding assessment, the source of the assessment, and link to the corresponding source data file.

**Table S10:** This table contains the pairwise chemical similarity values computed as the Tanimoto coefficient between ECFP4 fingerprints of the two chemicals. For each chemical, the corresponding PubChem identifier is provided.

**Table S11:** This table contains information on the biotransformation of 229 aquaculture chemicals. For each chemical, the table provides the PubChem identifier, Chemical Abstracts Service Registry Number (CASRN) and canonical SMILES of the aquaculture chemical, the corresponding successor chemical information such as SMILES and PubChem identiier (if available), the corresponding reaction name and SMILES, the enzyme involved in the reaction, the biosystem for the reaction, the type of transformation, and the source of the reaction.

**Table S12:** This table contains the edge list for the food web networks constructed using bioconcentration factor (BCF), acute toxicity, and chronic toxicity values. For each edge, the table provides the source node, target node, the corresponding edge weight (in ppm equivalents for chemical-species links), and the network type. Edges representing interactions between species were assigned an edge weight of 0.

**Table S13:** This table contains the node list for the food web networks constructed using bioconcentration factor (BCF), acute toxicity, and chronic toxicity values. For each node, the table provides the node label and type, betweenness centrality, and the corresponding network type.

## Supplementary Figure Captions

**Figure S1:** Sankey diagram depicting the flow of chemicals from their regulatory classes (left) to specific regulatory lists (right). The width of each connection is proportional to the number of unique chemicals. Values in parentheses (n) indicate the total unique chemical count for each node.

**Figure S2:** Chemical similarity network (CSN) of 339 discrete chemicals from the ReCAnt database. Edges represent Tanimoto coefficients computed using ECFP4 fingerprints. The figure illustrates 10 distinct groups, with each group comprising connected components with an identical number of constituent nodes. Maximum common substructures (MCS), constructed using RDKit, are displayed for each of the first five groups of connected components containing more than five nodes.

**Figure S3:** Directed biotransformation network consisting of 1510 nodes and 1380 edges. The network features 229 aquaculture chemicals (red nodes) and their 1281 associated transformation products (blue nodes).

**Figure S4:** Directed food web network constructed using BCF values to establish links between chemicals and prey species. The network comprises 348 edges connecting 120 nodes, of which 24 are chemicals (blue nodes), 47 are prey species (green nodes), 39 are predators (red nodes), and 10 species function as both predator and prey (orange nodes). Edge thickness represents the weight of the interaction between chemicals and prey.

