## Supplementary material for "A comprehensive resource on chemicals used in aquaculture and their ecotoxicity": Figure S

### <u>Supplementary Information</u>

### for

#### Supplementary Text

##### 1. Structural characterization of aquaculture chemicals

PubChem (https://pubchem.ncbi.nlm.nih.gov) is the world's largest openly-accessible online repository for chemical information, providing structural information in a standardized structure data file (SDF) format that allows for several structure-based analyses. Among the 690 curated chemicals, PubChem contained two-dimensional (2D) structural information for 339 chemicals, and these were therefore retrieved for further analyses. Next, Open Babel[1] (https://openbabel.org/) was utilized to generate structural identifiers such as the SMILES, InChI, and InChIKey. In addition to 2D structures, PubChem provided three-dimensional (3D) structural information only for 282 of 339 chemicals. Therefore, the Merck Molecular Force Field (MMFF94) method within Open Babel was used to generate the lowest-energy 3D structures from the 2D SDF files of the remaining 57 chemicals. Successful conversion was achieved for 48 compounds, while the remaining nine could not be processed because Open Babel was unable to generate their 3D structures. Open Babel was also utilized to convert 2D SDF files into 2D MOL and 2D MOL2 formats, and the 3D SDF files to 3D MOL, 3D MOL2, 3D PDB, and 3D PDBQT file formats. All of these files are made accessible through the online resource ReCAnt developed as part of this study.

Subsequently, ClassyFire[2] (http://classyfire.wishartlab.com) was used to classify the chemicals into hierarchical categories, namely, Kingdom, Class, and Subclass, for additional structural annotation. Furthermore, the RDKit module in Python was utilized to compute different physicochemical properties of these chemicals, such as molecular weight (in g/mol), logP, topological polar surface area (in Å$^2$), number of hydrogen bond donors/acceptors, carbon atoms, heavy atoms, heteroatoms, rotatable bonds, aliphatic rings, and aromatic rings. Finally, the 2D and 3D molecular descriptors were computed using PaDEL[3] (http://www.yapcwsoft.com/dd/padeldescriptor/), RDKit, and Pybel[4] for these 339 chemical structures. All of these files are made accessible through the online resource ReCAnt developed as part of this study.

#### 2. Assessment of regulatory coverage

To explore the extent of aquaculture-specific regulatory coverage of the compiled chemicals, the chemical lists provided by the Indian Coastal Aquaculture Authority (CAA) on banned chemicals for shrimp aquaculture[5], the United States Food and Drug Administration (US FDA) approved aquaculture drugs list (https://www.fda.gov/animal-veterinary/aquaculture/approved-aquaculture-drugs), and the Association of Southeast Asian Nations (ASEAN) Secretariat's 'Guidelines for the Use of Chemicals in Aquaculture and Measures to Eliminate the Use of Harmful Chemicals'[6], were relied upon. The Indian CAA-banned chemicals list consists of 29 chemicals, while the US FDA-approved aquaculture drugs list consists of 9 chemicals. The ASEAN guidelines provide 12 chemical lists that are categorized according to chemical classes and whether the fish are used for consumption or ornamentation. These lists comprise 97 chemicals spanning 8 countries, namely, Brunei Darussalam, Indonesia, Malaysia, Myanmar, Philippines, Singapore, Thailand, and Vietnam.

Furthermore, the European Chemical Agency's (ECHA) persistence, bioaccumulation, and toxicity (PBT) assessment list (https://echa.europa.eu/pbt) (last accessed on 22 November 2025), which includes PBT assessment for different compounds conducted by ECHA's PBT Expert Group, was retrieved. First, the dataset was restricted to assessments with a status of 'Concluded'. Entries classified as 'inconclusive', 'postponed', or undefined were then excluded, resulting in 155 compounds with PBT assessments. To expand the coverage of PBT-related assessments and to additionally incorporate mobility, the persistence, bioaccumulation, mobility, and toxicity (PBMT) datasets were curated from the NORMAN Suspect List

Exchange (https://www.norman-network.com/?q=suspect-list-exchange). Specifically, chemical lists titled UBAPMT[7] (Prioritised PMT/vPvM substances in the REACH registration database), EAWAGPMT[8] (PMT Suspect List from EAWAG), and UFZHSFPMT[9] (PMT Suspect List from UFZ and HSF), comprising 334, 1109, and 1031 compounds, respectively, were accessed. Moreover, the overlap of the curated 690 chemicals in our compilation with the Organisation for Economic Co-operation and Development (OECD) High Production Volume (OECD HPV) (https://hpvchemicals.oecd.org/ui/Default.aspx) database and the United States High Production Volume (US HPV) (https://comptox.epa.gov/dashboard/chemical-lists/EPAHPV) database was evaluated to capture information on production volumes. Additionally, 16 lists of substances of concern and substances in use from multiple jurisdictions, including India, the US, the European Union (EU), Japan, China, Australia, New Zealand, and Singapore, as well as from relevant regulatory bodies, were compiled based on lists in a previous work[10] that were relevant to this study. Together with the two HPV databases, a total of 18 lists were evaluated for overlap with the curated chemicals.

**3. Visualization of the chemical space**

The chemical similarity network (CSN)[11] for the 339 aquaculture-relevant chemicals that have 2D structures was constructed to visualize the chemical space through a network perspective. To this end, the Tanimoto coefficient[12] was employed as the similarity metric. Pairwise similarity coefficients were computed using ECFP4[13] fingerprints as implemented in the RDKit. Subsequently, a network was constructed, wherein the nodes are chemicals, and the edges are weighted by the corresponding Tanimoto coefficients. Thereafter, the edges with a Tanimoto coefficient > 0.5 were utilized to construct the CSN, which was visualized using Cytoscape version 3.10[14]. Next, to assess scaffold-level similarity among these chemicals, Bemis–Murcko scaffolds[15] were computed for each compound using RDKit. The frequency distribution of the resulting molecular scaffolds was then visualized using a scaffold cloud representation[16] generated with RDKit.

#### 4. Prediction of biotransformation, partitioning, and solubility of aquaculture chemicals

Chemicals can undergo transformation within environmental media, leading to the formation of transformation products (TPs),[17] or partition between different environmental compartments as governed by physicochemical processes.[18] To characterize these processes, the biotransformation pathways and environmental fate of aquaculture-relevant chemicals were investigated using a combination of computational tools and curated databases. First, the BioTransformer 3.0 tool[19] was used to predict potential TPs, with 2D SDF files provided as input. The biotransformation mode was set to environmental microbial degradation, which is informed by reaction rules and data from the EnviPath database,[20] and the predicted TPs were retrieved. As BioTransformer restricts its inputs to organic compounds with molecular masses below 1000 Da, the TP coverage was further enhanced by compiling additional TPs from databases available through the NORMAN Suspect List Exchange, namely HSDBTPS[21] (Transformation Products Extracted from HSDB Content in PubChem) and REFTPS[22] (Transformation Products and Reactions from Literature). To obtain an overview of the biotransformation landscape of aquaculture-related chemicals, a reaction network comprising all predicted transformation reactions was constructed and analyzed using NetworkX[23] and subsequently visualized in Cytoscape version 3.10[14].

Chemicals tend to partition among environmental media such as air, water, soil, and biota, and this behavior is commonly characterized using partition coefficients. A partition coefficient represents the ratio of a substance's concentration between two phases or environmental media at equilibrium under defined temperature and pressure conditions.[18] In this study, three widely used partition coefficients, namely $K_{ow}$ (octanol-water coefficient), $K_{aw}$

(air-water coefficient), and $K_{oa}$ (octanol-air coefficient), which together inform predictions of the chemical's environmental distribution and dominant partitioning behavior, were estimated.

Partition coefficients were estimated using the Estimation Programs Interface (EPI) Suite (https://episuite.dev/EpiWebSuite/#/), a screening-level tool developed by the US EPA that integrates multiple estimation programs to compute physicochemical and fate-related properties from a single input. The CASRNs of aquaculture-relevant chemicals were used as input to estimate the three partition coefficients, as well as the BCF (L/kg wet-weight) and bioaccumulation factor (BAF; L/kg wet-weight). In addition, EPI Suite provides quantitative structure–activity relationship (QSAR)-based toxicity predictions via the EPA Ecological Structure Activity Relationships (ECOSAR) Predictive Model (https://www.epa.gov/tsca-screening-tools/ecological-structure-activity-relationships-ecosar-predictive-model), which estimates concentration thresholds for multiple toxicity endpoints across organisms and exposure durations, and the QSAR classes based on different structural features of the chemical. Here, ECOSAR predictions were restricted to aquatic organisms and, as model-derived estimates, were excluded from downstream analyses and retained only for preliminary hazard assessment.

Further, the solubility for the aquaculture-relevant chemicals was retrieved from AqSolDB,[24] which is a curated database on aqueous solubility for a diverse set of chemicals. AqSolDB reports the logarithm of solubility (LogS), along with the corresponding standard deviation across values from primary data sources. Based on the classification provided by AqSolDB, chemicals were classified as highly soluble (LogS is 0 or higher), soluble (0 to -2), slightly soluble (-2 to -4), and insoluble (-4 or lower).[24] The data on biotransformation, chemical fate, toxicity predictions, and solubility are accessible through the online resource ReCAnt developed as part of this study.

#### 5. Analyses of trophic transfer potential of aquaculture chemicals through food webs

Despite limited direct environmental exposure, aquaculture chemicals can affect species at higher trophic levels through dietary uptake resulting from the consumption of contaminated organisms at lower trophic levels.[25] Furthermore, the transfer of chemicals into food webs can alter predator–prey dynamics by disproportionately affecting the abundance, fitness, or survival of either prey or predator species, thereby reshaping trophic interactions within the ecosystem.[26,27] To investigate how aquaculture chemicals may enter food webs associated with aquaculture-relevant species and exert acute or chronic toxic effects or undergo bioaccumulation, the food web interaction data was retrieved from the Global Biotic Interactions (GloBI) (https://www.globalbioticinteractions.org),[28] which is one of the largest databases on species interaction data curated from open datasets. Specifically, interactions in which aquaculture-relevant species function as predators were extracted and integrated with prey-level bioconcentration factor (BCF) and toxicity data from ECOTOX. For this analysis, it was assumed that aquaculture-relevant species are not co-cultured with their natural predators, and that although prey species may not all co-occur in the same system, they represent potential dietary sources for predator species. This integration enabled assessment of chemical exposure at lower trophic levels and the potential for transfer to aquaculture species through trophic interactions.

First, the GloBI database was queried on 17 November 2025 through the GloBI Web API, using a compiled list of 81 aquaculture-relevant species, and the obtained results were parsed using Python. Interactions were restricted to the interaction types ‘eats’ and ‘preysOn’, with aquaculture-relevant species specified as the source taxa. This query returned food-web interactions involving 77 aquaculture-relevant species connected to 4128 taxa through a total of 12464 interactions. Next, acute toxicity, chronic toxicity, and BCF data from ECOTOX were integrated at the prey level of the food web using the workflow described in Section 2.3 of the main text. This integration resulted in three directed networks, in which edge weights represented BCFs or toxicity concentration values.

Network analysis was performed using the NetworkX[23] module in Python to quantify connectivity between chemicals and predator species. It should be noted that for this analysis, self-loops signifying intraspecific predation were removed, as there is no trophic level change in such cases. For the BCF-appended food web network, the existence of any path between a chemical and predator node was used as a proxy for bioaccumulation potential through trophic transfer. For toxicity-appended networks (acute and chronic), path existence indicated potential dietary exposure to chemicals causing toxic effects at prey levels. For each network, the number of predator species connected to each chemical and the number of chemicals connected to each predator species were calculated to identify chemicals with high potential for trophic transfer and predator species with high exposure vulnerability. Betweenness centrality was calculated for all prey species in each network to quantify how often each species lies on the shortest paths between chemicals and predators. This calculation treated the network as unweighted, as connections between predator and prey species carried an edge weight of 0. Higher centrality values potentially indicate species that serve as important bridges in chemical exposure pathways to higher trophic levels. Networks were visualized in Cytoscape version 3.10.[14]

**6. Web interface and database management system**

To enable access to the wide range of data generated in this study, an online resource, namely, Resource on Chemicals used in Aquaculture and their Ecotoxicity (ReCAnt), was developed and is made accessible for academic research at: https://cb.imsc.res.in/recant/. For each chemical, ReCAnt provides comprehensive information, including downloadable structure files, molecular descriptors, physicochemical properties, curated therapeutic potential data, curated toxic effect data, CTD associations, and ECOTOX-derived toxicity and BCF data.

In addition, ReCAnt integrates chemical screening information, including predicted biotransformation products and partition coefficients, and provides network-based visualizations of biotransformation relationships depicting each chemical and its first-neighbor transformation products. Food web-associated data are presented through predator-centric network visualizations, in which aquaculture-relevant species serve as predators, highlighting prey species that may contribute to trophic transfer via toxicity or bioaccumulation.

The compiled data is stored in a MariaDB (https://mariadb.org/) database and retrieved using Structured Query Language (SQL). The web interface was developed using HTML, CSS, Bootstrap 5 (https://getbootstrap.com/docs/5.0/), jQuery (https://jquery.com/), and PHP (http://php.net/), and hosted on an Apache webserver (https://httpd.apache.org/) running on the Debian 9.4 operating system. Network visualizations are rendered using the Cytoscape.js graph library (http://js.cytoscape.org/).[29]

#### Supplementary Figures

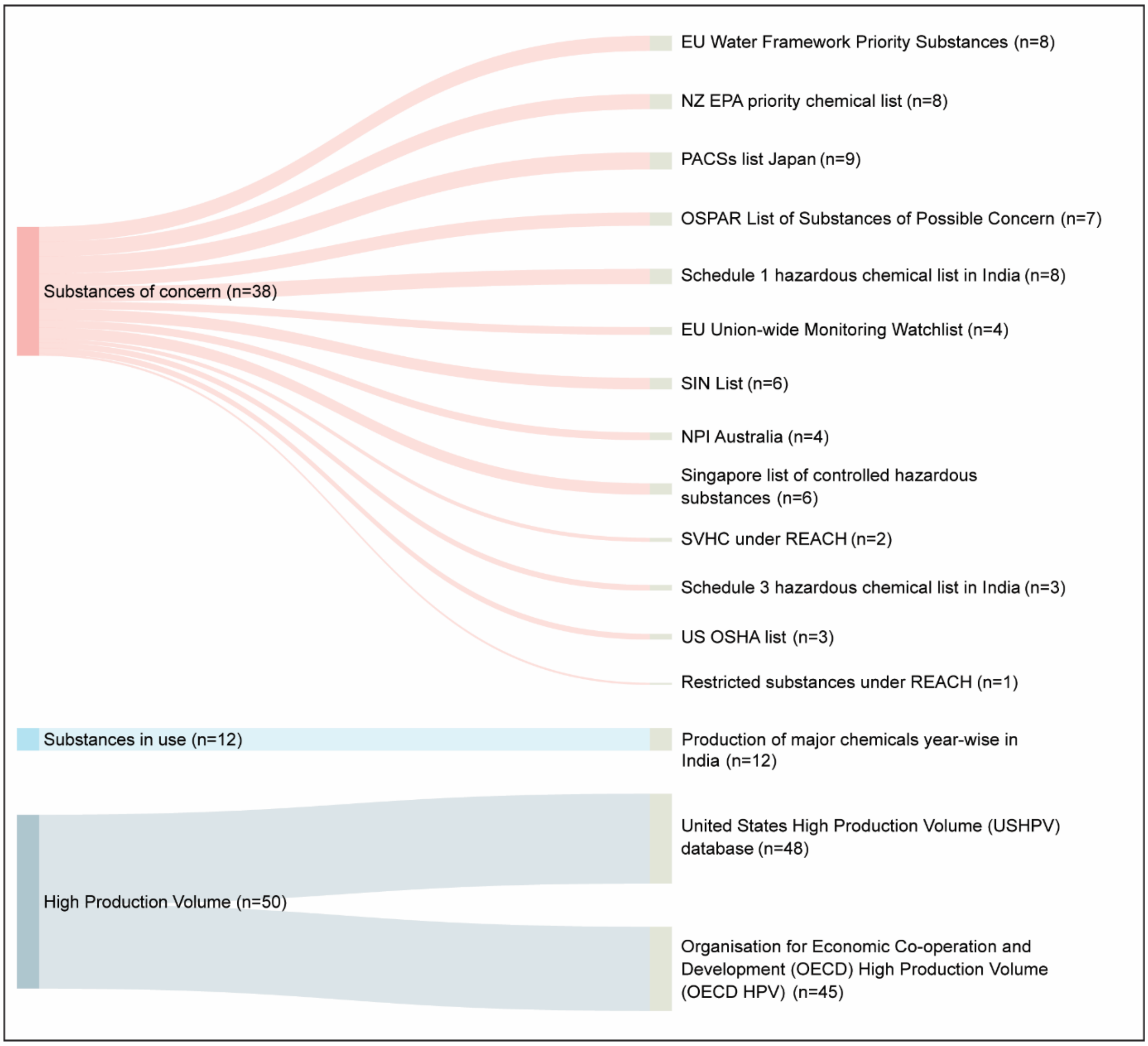


**Figure S1:** Sankey diagram depicting the flow of chemicals from their regulatory classes (left) to specific regulatory lists (right). The width of each connection is proportional to the number of unique chemicals. Values in parentheses (n) indicate the total unique chemical count for each node.

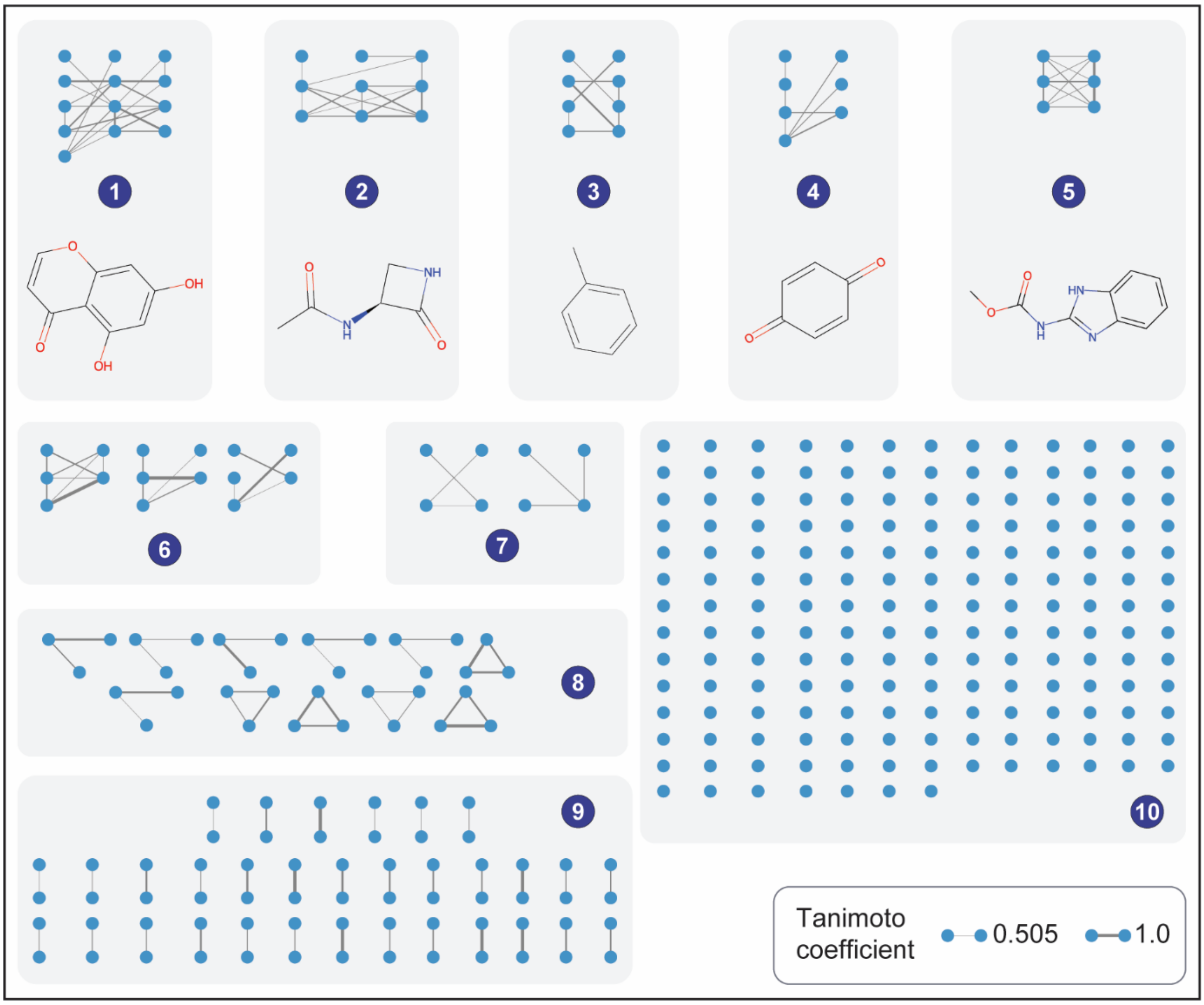


**Figure S2:** Chemical similarity network (CSN) of 339 discrete chemicals in the ReCAnt database. Edges represent Tanimoto coefficients computed using ECFP4 fingerprints. The figure illustrates 10 distinct groups, with each group comprising connected components with an identical number of constituent nodes. Maximum common substructures (MCS), constructed using RDKit, are displayed for each of the first five groups of connected components containing more than five nodes.

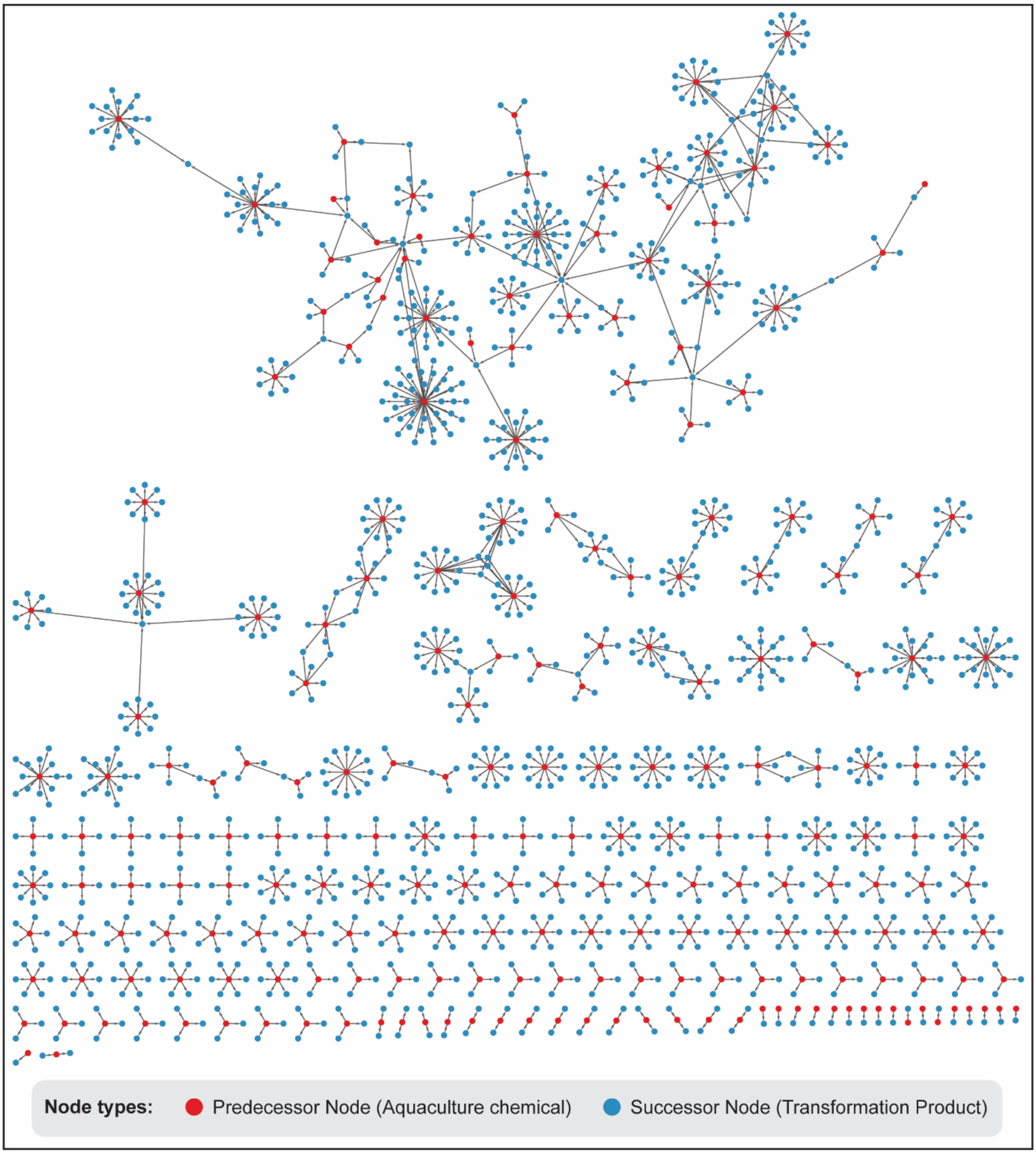


**Figure S3**: Directed biotransformation network consisting of 1510 nodes and 1380 edges. The network features 229 aquaculture chemicals (red nodes) and their 1281 associated transformation products (blue nodes).

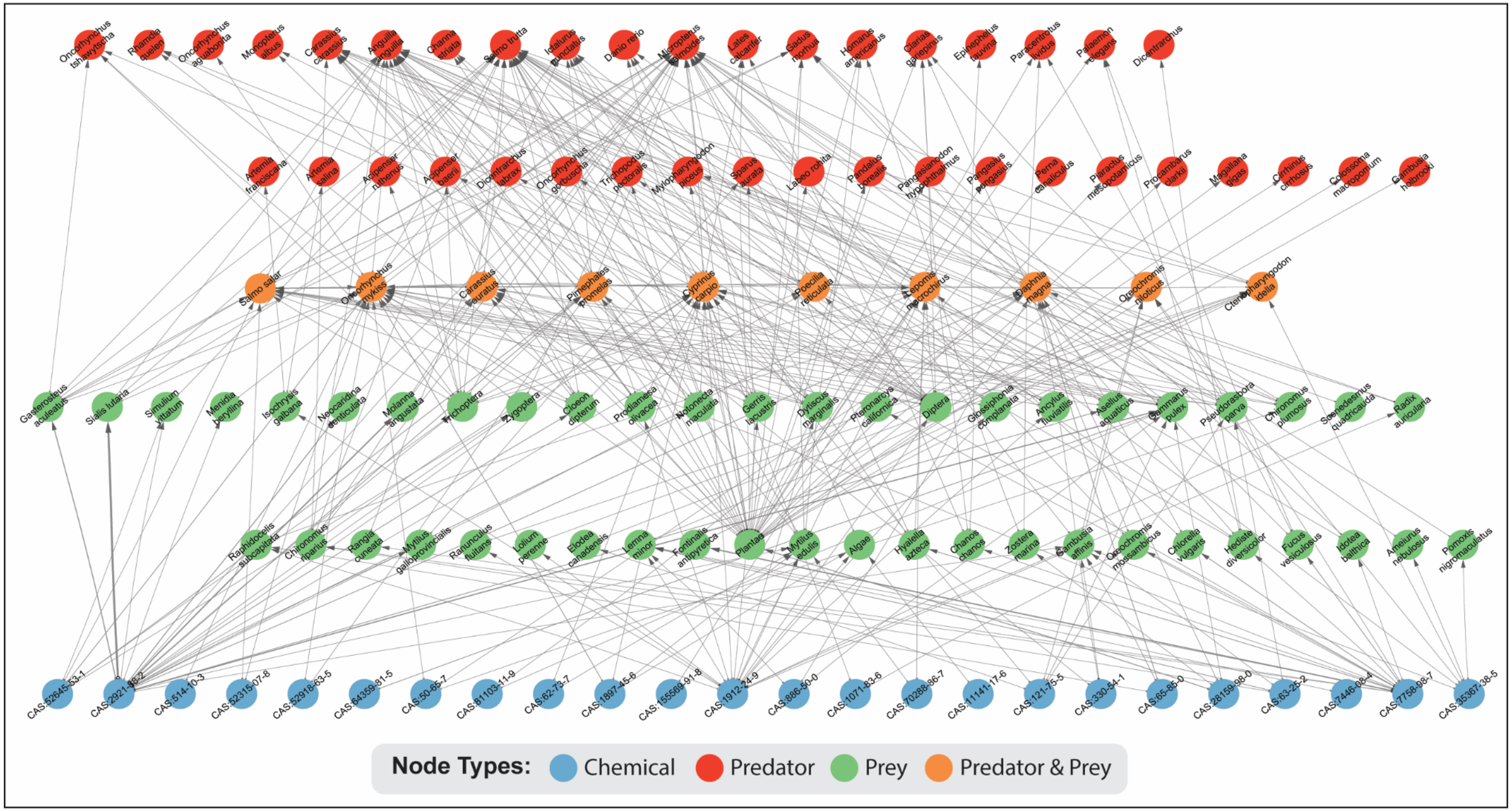


**Figure S4**: Directed food web network constructed using BCF values to establish links between chemicals and prey species. The network comprises 348 edges connecting 120 nodes, of which 24 are chemicals (blue nodes), 47 are prey species (green nodes), 39 are predators (red nodes), and 10 species function as both predator and prey (orange nodes). Edge thickness represents the weight of the interaction between chemicals and prey.
